# A Beta-to-Alpha seasonal coronavirus antibody signature identifies high and low responders to SARS-CoV-2 vaccination

**DOI:** 10.64898/2026.11.15.766755

**Authors:** Stephanie Longet, Caolann Brady, Ivan Tomic, Tom Tipton, Salma Bessalah, Eleanor Barnes, Susanna Dunachie, Christopher JA Duncan, Paul Klenerman, Alex Richter, James ED Thaventhiran, Lance Turtle, Thushan I de Silva, Adriana Tomic, Miles W Carroll

**Affiliations:** Centre for Human Genetics and Pandemic Sciences Institute, Nuffield Department of Medicine, University of Oxford, Oxford, United Kingdom; Pandemic Sciences Institute, University of Oxford, Oxford, United Kingdom; International Center for Infectiology Research (CIRI), Team GIMAP, Claude Bernard Lyon 1 University, Inserm, U1111, CNRS, UMR530, Saint-Etienne, France; Atomic Lab, National Emerging Infectious Diseases Laboratories, Boston University, 620 Albany St, Boston, MA 02118, United States; Department of Virology, Immunology & Microbiology, Boston University Medical School; Boston, MA, US; Biomedical Engineering, Boston University, College of Engineering; Boston, MA, US; Nuffield Department of Clinical Medicine, Peter Medawar Building for Pathogen Research, University of Oxford, Oxford, United Kingdom; Oxford University Hospitals NHS Foundation Trust, Oxford, United Kingdom; Nuffield Department of Clinical Medicine, Oxford Centre For Global Health Research, University of Oxford, Oxford, United Kingdom; Mahidol-Oxford Tropical Medicine Research Unit, Mahidol University, Bangkok, Thailand; Translational and Clinical Research Institute Immunity and Inflammation Theme, Newcastle University, Newcastle, United Kingdom; Department of Infection and Tropical Medicine, Newcastle upon Tyne Hospitals NHS Foundation Trust, Newcastle, United Kingdom; Translational Gastroenterology Unit, University of Oxford, Oxford, United Kingdom; NIHR Oxford Biomedical Research Centre, University of Oxford, Oxford, United Kingdom; Institute of Cancer and Genomic Science, College of Medical and Dental Science, University of Birmingham, Birmingham, United Kingdom; University Hospitals Birmingham NHS Foundation Trust, Birmingham, United Kingdom; MRC Toxicology Unit, University of Cambridge, Cambridge, UK; NIHR Health Protection Research Unit in Emerging and Zoonotic Infections, Institute of Infection, Veterinary and Ecological Sciences, University of Liverpool, Liverpool, United Kingdom; Liverpool University Hospitals NHS Foundation Trust, Liverpool, United Kingdom; Department of Infection, Immunity and Cardiovascular Disease, University of Sheffield, Sheffield, United Kingdom; Sheffield Teaching Hospitals NHS Foundation Trust, Sheffield, United Kingdom

## Abstract

Pre-existing immunity to endemic seasonal human coronaviruses may shape subsequent responses to SARS-CoV-2 vaccination. We longitudinally profiled 775 healthcare workers using a systems immunology machine learning framework and found that a recursive feature elimination cross-validation model classified individuals in the top and bottom quartiles of SARS-CoV-2 Spike IgG responses (AUC=0.72, empirical P<0.001). This predictive signature was prominently driven by a baseline bias toward Beta-coronavirus relative to Alpha-coronavirus Spike reactivity.

## Main text

The Severe Acute Respiratory Syndrome Coronavirus 2 (SARS-CoV-2) trimeric spike (S) protein, composed of 1273 amino acids and divided into two subunits (S1 and S2), has been shown to be the main target of antigen-specific antibodies essential for protection after SARS-CoV-2 infection and COVID-19 vaccination ^1,2^. A comparison of S protein sequences using the NCBI protein BLAST tool showed a variable degree of amino-acid sequence identity between the S sequences of SARS-CoV-2 (QOS45029.1) and all human coronaviruses (hCoVs) including the betacoronaviruses OC43-CoV (UOP57234.1) (36.32%), HKU1-CoV (BDB58097.1) (36.79%), SARS-CoV-1 (AAR86788.1) (75.88%) and MERS-CoV (QGV13484.1) (35.19%), as well as the alphacoronaviruses NL63-CoV (XEQ84071.1) (30.32%) and 229E-CoV (XEQ84004.1) (30.94%)^3^. This sequence and structural relatedness may lead to the presence or expansion of circulating cross-reactive antibodies. This was highlighted by the presence of pre-existing humoral immunity against SARS-CoV-2 in unexposed individuals which was detected early in the SARS-CoV-2 pandemic ^4^. An enhancement of S-specific IgG responses to the hCoVs, and particularly the betacoronaviruses OC43 and HKU1, were also observed in COVID-19 convalescent individuals ^5–7^. In addition, the impact of cross-reactive antibodies on COVID-19 disease was analysed in several studies but it remains unclear whether and how they influence clinical outcome. A range of human and preclinical trials observed that hCoV cross-reactive immunity could have a negative impact on COVID-19 clinical outcomes ^8–11^, even though there has not been any evidence of antibody-dependent enhancement in SARS-CoV-2 infection or COVID-19 vaccination ^12^. Some studies demonstrated that hCoV cross-reactive immunity could protect against COVID-19 ^13,14^, while others did not observe any clinical benefit on COVID-19 outcomes ^7,15–18^. Antibody cross-reactivity was also evaluated following COVID-19 vaccination. It was observed in our studies and studies from other research teams that mRNA COVID-19 vaccination boosted the serum IgG reactivity to S of HKU1 and OC43 ^19–25^. Thus, pre-existing immunity to hCoVs shapes subsequent SARS-CoV-2 immune responses. However, whether baseline hCoV S-specific IgG profiles can predict SARS-CoV-2 vaccine-induced S-specific IgG responses and their longitudinal trajectories across diverse populations remains unclear.

Machine learning (ML) has been used to unravel the complexities of vaccine-specific immune responses and to predict immune signatures after vaccination. Sequential Iterative Modelling OverNight (SIMON) is a novel automated ML system which compares several hundred algorithms to identify the algorithm that best models a high dimensional dataset ^26,27^. This ML system has been used to determine immune signatures of Influenza vaccine responses using datasets from five clinical trials ^26^. In addition, it successfully predicted Typhoid and Meningococcus B vaccine immune responses ^27,28^. In the context of SARS-CoV-2 infection, SIMON identified early humoral and cellular immune predictors of long-term immunity in UK healthcare workers ^6^, immune signatures linked with COVID-19 severity^29^ and clinical and viral protection following SARS-CoV-2 challenge in a cohort of immunised non-human primates ^11^.

In this study, we longitudinally profiled 775 healthcare workers (HCW) using a systems immunology machine learning framework, combined with quartile-based response classification and stringent correction for broad heterologous coronavirus reactivity. SARS-CoV-2 previously infected and naive HCW were vaccinated with COVID-19 mRNA or ChAdOx-based vaccines approved in the UK between end of 2020 and 2022, and their blood was collected at different timepoints before vaccination (baseline) and after one, two and three doses to measure SARS-CoV-2, SARS-CoV-1, MERS-CoV, HKU1, OC43, NL63, 229E S-specific IgG responses. Potential SARS-CoV-2 exposures were monitored using SARS-CoV-2 Nucleocapsid (N)-specific IgG responses. We found that N-specific IgG waned independently of S-specific IgG responses (Figure 1), which ruled out intercurrent SARS-CoV-2 infection or exposure as a cause of S-specific IgG boosting. Rather, vaccination alone was responsible for S-specific IgG boosting.

**Figure 1:**
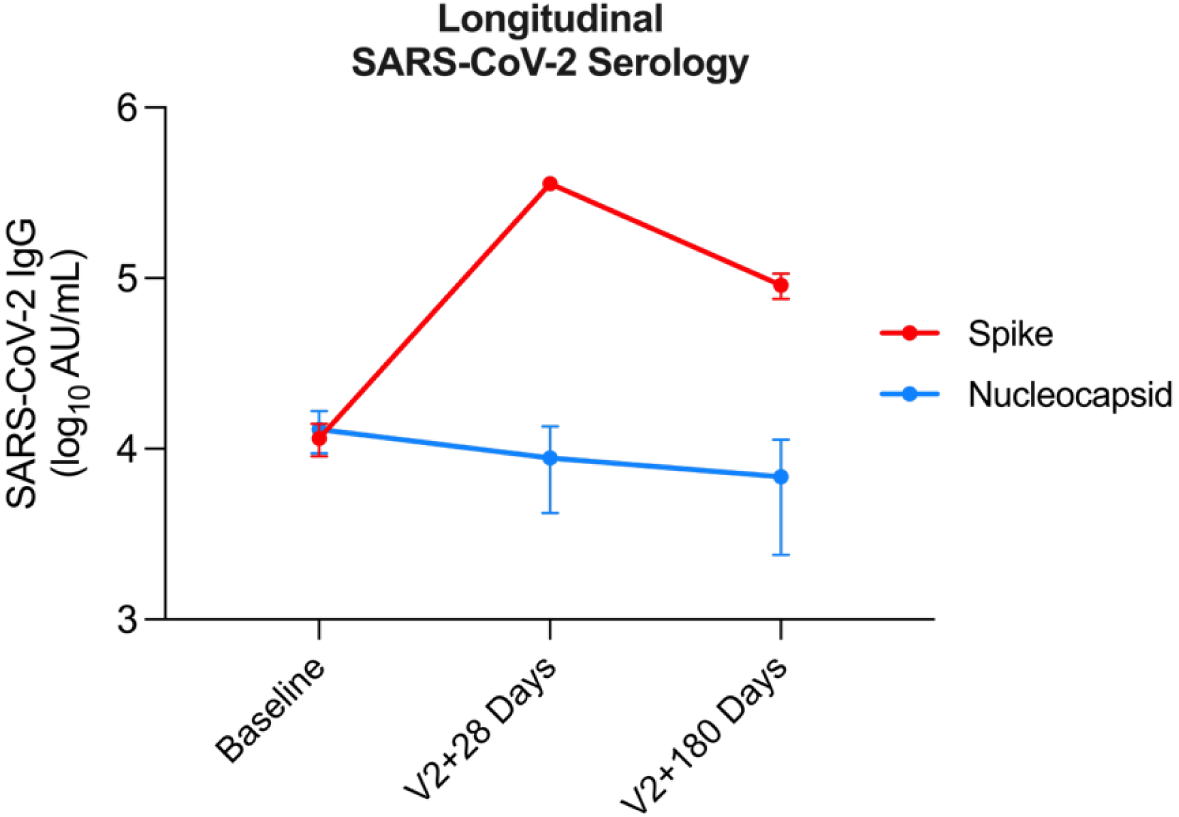
Nucleocapsid IgG Waning as an Internal Biological Control. Longitudinal trajectories of SARS-CoV-2 Nucleocapsid (N)- and Spike (S)-specific IgG titers (log_10_AU/mL) in previously infected healthcare workers at baseline (pre-vaccination), 28 days post-second dose (v2+28), and 180 days post-second dose (v2+180). Continuous waning of N-specific IgG confirms the antigen-specificity of vaccine-induced Spike boosting and decay kinetics, ruling out intercurrent SARS-CoV-2 re-exposure during follow-up. Datapoints represent the mean with the bars representing the 95% confidence interval.

In this UK HCW cohort, robust true memory against MERS and SARS-CoV-1 is biologically improbable. High binding to these antigens was therefore interpreted as reflecting non-specific assay polyreactivity or low-affinity broad heterologous coronavirus cross-reactivity. Therefore, to separate the signal of endemic seasonal coronavirus memory from a signal representative of broad heterologous coronavirus reactivity, a conservative correction for broad heterologous coronavirus reactivity was established to regress out the component of hCoV S-specific IgG signal associated with MERS-CoV and SARS-CoV-1 S-specific IgG binding. This enabled us to include cross-reactivity-corrected hCoV S-specific IgG response as a possible baseline serological feature associated with SARS-CoV-2 S-specific IgG outcome post-vaccination. By regressing this background reactivity out, this approach isolates the seasonal hCoV S-specific signal from a non-specific cross-reactive binding background signal.

Additional baseline hCoV serological features included baseline log10-transformed coronavirus serology, Beta/Alpha coronavirus immune-bias score and cumulative hCoV breadth. Beta/Alpha coronavirus immune-bias score is a baseline serology-derived metric contrasting corrected S-specific IgG reactivity to seasonal Beta-coronaviruses, OC43 and HKU1, with corrected S-specific IgG reactivity to seasonal Alpha-coronaviruses, 229E and NL63. This metric was designed to capture whether pre-existing coronavirus immunity was relatively enriched for OC43/HKU1-like Beta-coronavirus reactivity, which may be more structurally relevant to SARS-CoV-2 than Alpha-coronavirus reactivity.

With this dataset, we aimed to address two questions, namely, does baseline hCoV immunity infer SARS-CoV-2 vaccine serological response 1) longevity, and/or 2) magnitude?

In cross-sectional ordinary least-squares modeling of participant-specific antibody decay between 28 and 180 days post-second dose, baseline endemic hCoV titers did not significantly predict antibody waning rate (R^2^ = 0.13, p = 0.091). Prior SARS-CoV-2 infection, did however, significantly attenuated the rate of antibody waning (β = −0.287, P = 0.05), demonstrating that hybrid immunity enhances the long-term durability of vaccine-induced Spike IgG compared to infection-naive individuals.

To determine whether baseline hCoV serological features were associated with SARS-CoV-2 vaccine serological response, we utilised SIMON. We found that standard continuous ML on the full cohort did not robustly predict individual SARS-CoV-2 S-specific IgG response magnitudes across the full response distribution. We therefore used a quartile-based classification strategy to focus on the upper and lower ends of the vaccine-response distribution. Individuals in the upper quartile were compared with individuals in the lower quartile, while the middle two quartiles were not included in the binary classification analysis. This enabled us to test whether baseline hCoV-derived features could discriminate the extremes of vaccine responsiveness.

A Recursive Feature Elimination Cross-Validation (RFECV) model, with an area under the receiver-operating characteristic curve (AUC) of 0.72, discriminated participants with the highest and lowest quartiles of vaccine-induced SARS-CoV-2 S-specific IgG responses (Figure 2). To assess whether this performance exceeded that expected by chance, we performed 1,000 label-permutation analyses, which yielded an empirical P value <0.001 (Figure 3).

**Figure 2:**
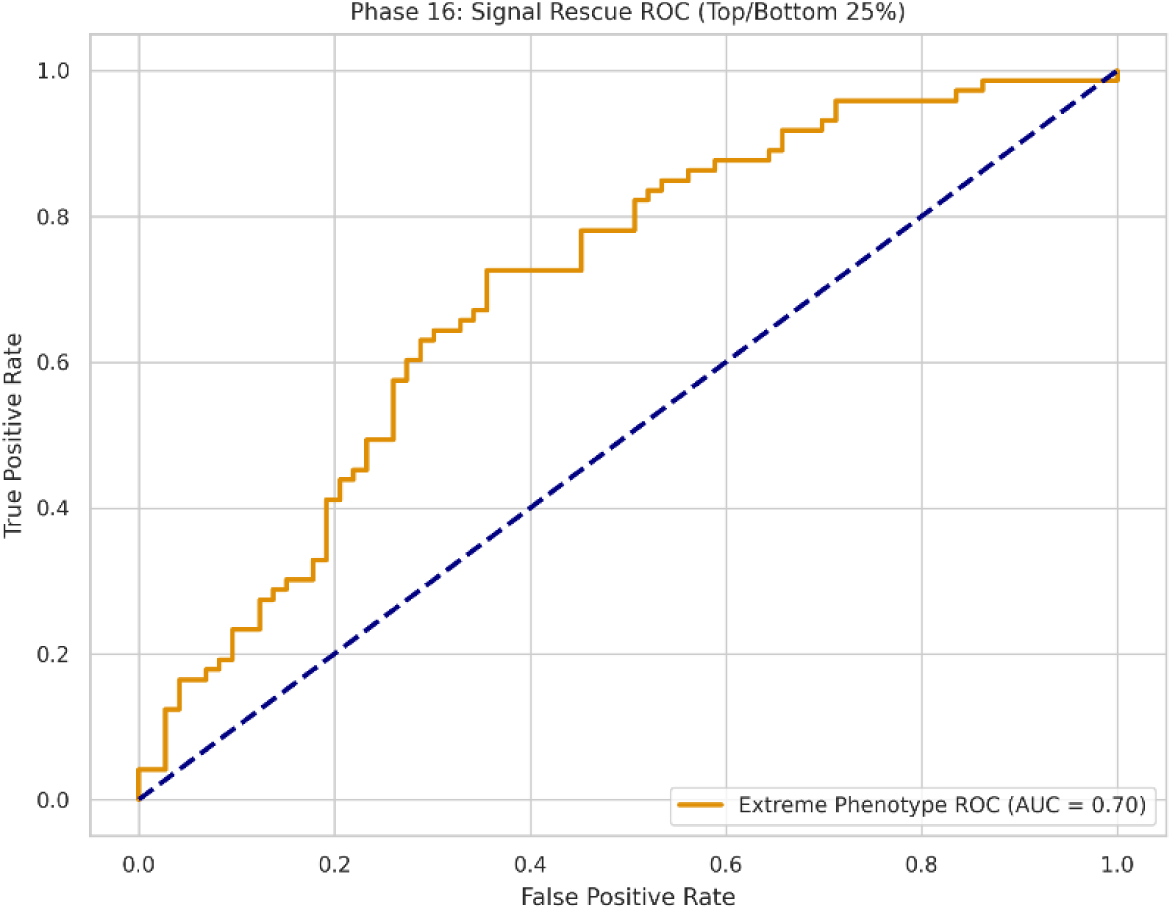
Receiver Operating Characteristic (ROC) Curve of the Extreme Phenotyping Predictive Model. The ROC curve illustrates model classification performance on the holdout evaluation set (AUC = 0.70), discriminating individuals in the upper and lower quartiles of SARS-CoV-2 Spike IgG response. Mean 5-fold cross-validation performance across the training cohort achieved an AUC of 0.72 (empirical P < 0.001 by 1,000-permutation testing).

**Figure 3:**
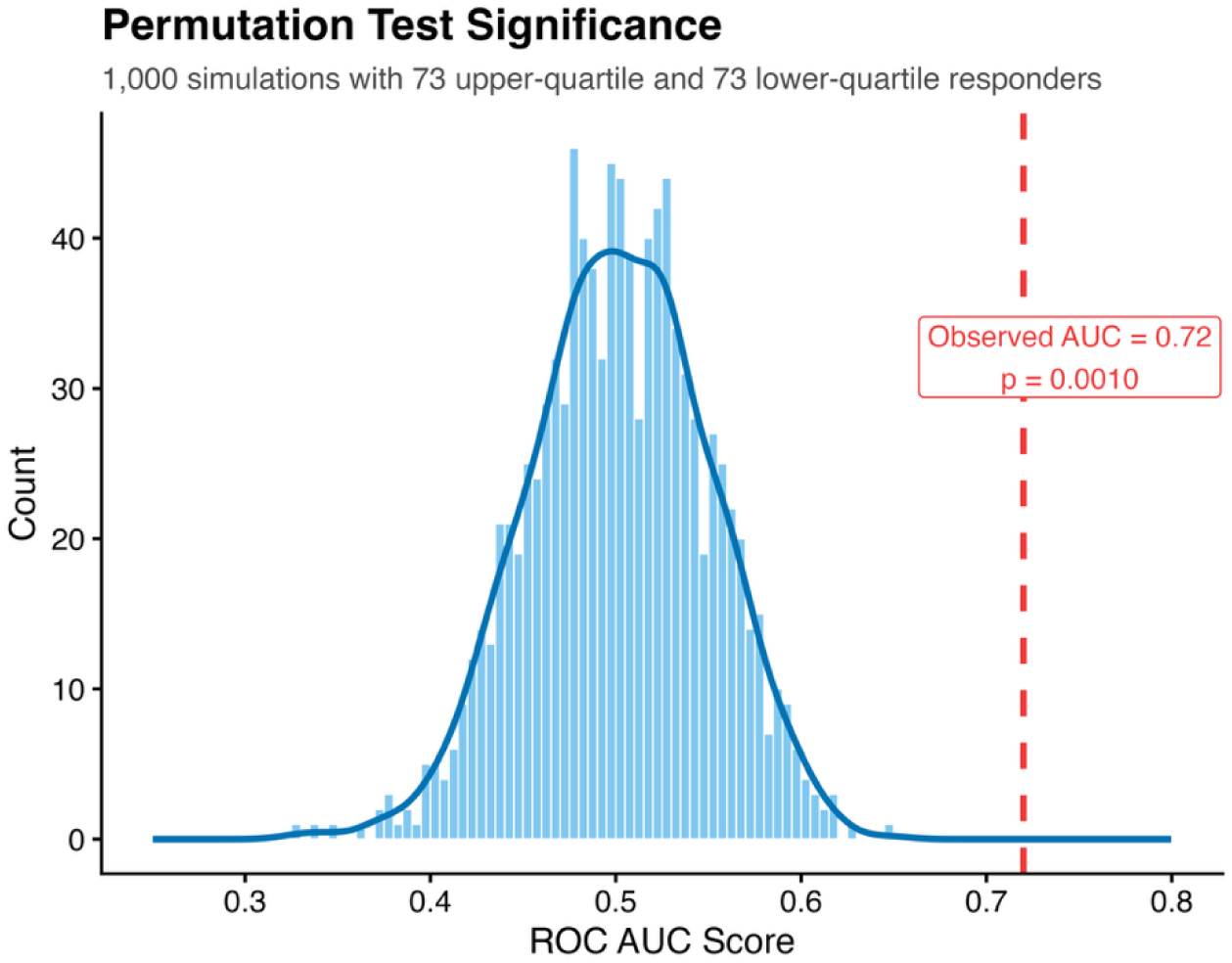
The Permutation Null-Distribution. A thousand permutations with scrambled labels were run yielding an empirical p < 0.001.

The model isolated an optimal eight-feature predictive signature comprising host demographics (age, sex, and prior SARS-CoV-2 infection status), cross-reactivity corrected seasonal titers (229E, HKU1, and NL63), and the two engineered systems features: the Beta/Alpha coronavirus immune-bias score and cumulative hCoV breadth. It is important to note that, Beta/Alpha immune-bias score and the cumulative HCoV breadth score captured the variance of cross-reactivity corrected OC43 S-specific IgG titers such that it was discarded by the model and therefore not featured in the predictive signature.

Interestingly, the Beta/Alpha coronavirus immune-bias score prominently drove this predictive signature. Thus, a pre-existing Beta-coronavirus-skewed antibody profile was associated with the upper and lower ranges of SARS-CoV-2 vaccine-induced humoral immunity. Consistent with shared or cross-reactive S epitopes between SARS-CoV-2 and endemic Beta-coronaviruses, OC43 S-specific IgG binding was significantly elevated after COVID-19 vaccination in this study (Figure 4), in agreement with previous reports^19–25^.

**Figure 4:**
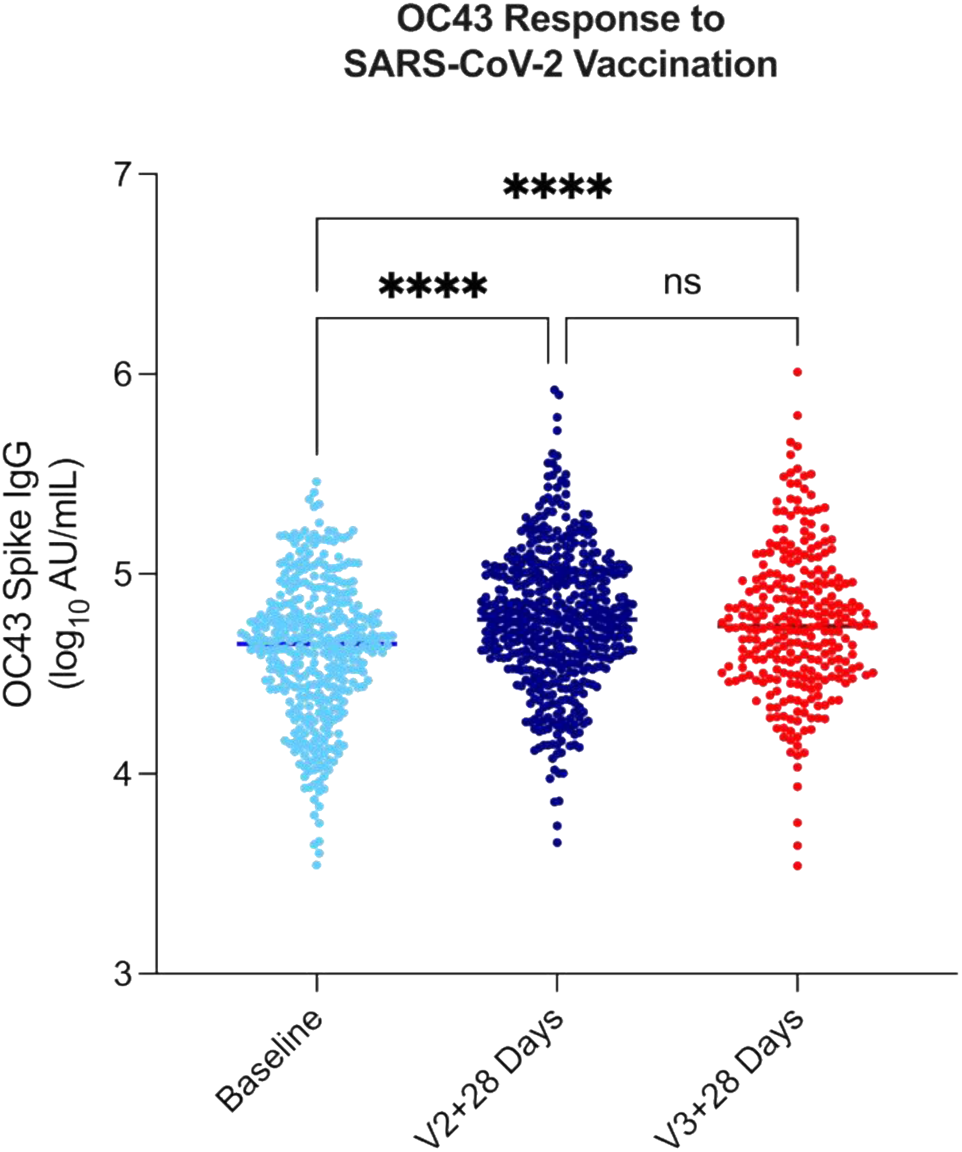
OC43 S-specific IgG Recall responses. OC43 S-specific IgG binding is significantly increased after COVID-19 vaccination, proving structural epitope sharing between SARS-CoV-2 and endemic Beta-coronaviruses. Baseline: OC43 S-specific IgG titers (log10) before vaccination. V2plus28: OC43 S-specific IgG titers (log10) 28 days after the second dose. V3plus28: OC43 S-specific IgG titers (log10) 28 days after the third dose. One-way ANOVA with Holm-Šídák’s multiple comparisons test performed with adjusted p-value displayed, p < 0.0001. Datapoints represent each participant. Median represented by a line.

Thus, our predictive models support the conclusion that pre-existing Beta-coronavirus-skewed baseline serology, quantified by a Beta/Alpha coronavirus immune-bias score, helps discriminate individuals at the upper and lower ends of SARS-CoV-2 vaccine-induced humoral immunity. Although generalized priming for coronavirus recognition is plausible, our data are consistent with a model in which structural similarity among Beta-coronaviruses contributes to recall or expansion of cross-reactive IgG specificities. The selective enhancement of Beta-coronavirus S-specific IgG binding after COVID-19 vaccination aligns with the conserved 3D structural similarity shared between SARS-CoV-2, OC43, and HKU1, particularly within the S2 Spike stem, which is less conserved among the divergent Alpha-coronaviruses.

While this serological study cannot definitively differentiate between the reactivation of pre-existing OC43 clones and the generation of highly cross-reactive *de novo* SARS-CoV-2 clones, as this would require cellular mechanistic studies, single B-cell or B-cell receptor lineage analyses, and/or bone marrow B-cell profiling, the predictive power of the Beta/Alpha coronavirus immune-bias score suggests that shared Beta-coronavirus antigenic features contribute to the observed serological pattern. Seasonal hCoV immunity, however, was not associated with vaccine-induced serological response durability up to 180 days following the second dose. Rather a prior SARS-CoV-2 infection was associated with humoral response longevity. Ultimately, these findings indicate that the capacity to mount a robust humoral response to a novel pandemic antigen may be influenced by structural overlap with the host’s pre-existing endemic memory B-cell repertoire, and humoral response durability can be maximised with hybrid immunity.

## Methods

### Study design

This study is part of the Protective Immunity from T-cells in Healthcare Workers (HCW) (PITCH) consortium, which is a prospective, observational, cohort study, with HCWs recruited from across six NHS hospitals in the UK (Birmingham, Liverpool, Newcastle, Oxford, Sheffield and Cambridge).

PITCH is a sub-study of the SIREN study, which was approved by the Berkshire Research Ethics Committee, Health Research 250 Authority (IRAS ID 284460, REC reference 20/SC/0230), with PITCH recognised as a sub-study on 2 December 2020. SIREN is registered with ISRCTN (Trial ID:252 ISRCTN11041050). The study was conducted in compliance with all relevant ethical regulations for work with human participants, and according to the principles of the Declaration of Helsinki (2008) and the International Conference on Harmonization (ICH) Good Clinical Practice (GCP) guidelines. Written informed consent was obtained for all participants enrolled in the study. Eligible participants were adults aged 18 or over, and currently working as a HCW, including allied support and laboratory staff, or were volunteers linked to the hospital.

Participants were sampled between December 2020 and February 2022, with the majority of the sampling completed before the omicron BA.1 variant emerged in the UK. Participants had received one, two or three doses of BNT162b2 (Pfizer/BioNTech), AZD1222 (Oxford/AstraZeneca) or mRNA-1273 (Moderna). The doses were administered with a dosing interval from 19 to 105 days. Participants underwent phlebotomy for assessment of immune responses, 7 days, 14 days, 28 days after the first dose and 7, 14, 28 and 180 days after the second dose of vaccine, and 28, 180 days after the third vaccine dose.

Information including BNT162b2 and AZD1222 vaccination dates, date of any SARS-CoV-2 pre-infection defined by a positive PCR test and/or detection of antibodies to S or N proteins, age and sex of participant was recorded. Key information on demographics of the cohort included in this study is shown in **Table 1**. Participants were divided into SARS-CoV-2 previously-infected and naïve individuals before vaccination. Participants were considered to be SARS-CoV-2 exposed if they had ever been PCR or lateral flow device positive for SARS-CoV-2, irrespective of symptoms. In addition, participants were considered exposed to SARS-CoV-2 if they seroconverted with N antibody on the Meso Scale Discovery (MSD) assay.

**Table 1:** Demographic characteristics of participants.

|  | **Naive** | **Previously-infected** |
|---|---|---|
| **N** | 449 | 326 |
| **Age in years (IQR)** | 41 (29-51) | 41 (30-51) |
| **Female (%)** | 301 (67%) | 255 (78%) |
| **Male (%)** | 148 (33%) | 71 (22%) |

### Meso Scale Discovery (MSD) IgG binding assay

Serum IgG responses to Wuhan SARS-CoV-2, SARS-CoV-1, MERS-CoV, HKU1, OC43, NL63, 229E S antigens were measured using multiplex MSD immunoassay: The V-PLEX COVID-19 Coronavirus Panel 3 (IgG) (cat# K15399U) from Meso Scale Discovery, Rockville, MD USA. A MULTI-SPOT® 96-well, 10 spot plate was coated with three Wuhan SARS CoV-2 antigens (S, Receptor-Binding Domain (RBD), N), SARS-CoV-1 and MERS-CoV S trimers, S proteins from OC43, HKU1, 229E and NL63, and bovine serum albumin (negative control). Multiplex MSD assays were performed as per the manufacturer’s instructions. To measure IgG antibodies, 96-well plates were blocked with MSD Blocker A for 30 minutes. Following washing, samples diluted 1:1,000-40,000 in diluent buffer, MSD standard and undiluted internal MSD controls, were added to the wells. After 2-hour incubation and a washing step, detection antibody (MSD SULFO-TAG™ anti-human IgG antibody, 1/200) was added. Following washing, MSD GOLD™ read buffer B was added and plates were read using a MESO® SECTOR S 600 reader. The standard curve was established by fitting the signals from the standard using a 4-parameter logistic model. Concentrations of samples were determined from the electrochemiluminescence signals by back-fitting to the standard curve and multiplying by the dilution factor. Concentrations are expressed in Arbitrary Units/ml (AU/ml).

N seroconversion was defined as an N antibody level over the cut-off threshold of 3874 previously defined using 103 pre-pandemic samples + 3 Standard Deviations ^20^, and at least a 2-fold increase over the baseline value. Participants who did not meet any of these criteria were considered to be infection-naïve. Most data has been previously published ^20,21,30^.

### Pre-processing of serological data

Raw luminescence values were pre-processed before feature engineering. Values below the analytical floor were set to 1, after which all serological measurements were log10-transformed:

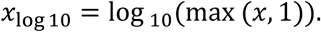

This transformation was applied to reduce skewness in antibody-binding measurements and to place low or undetectable responses at zero on the log10 scale.

### Stringent cross-reactivity penalization

To isolate endemic seasonal coronavirus memory from broad heterologous coronavirus reactivity, we implemented a stringent cross-reactivity penalization strategy. For each participant, a conservative background-reactivity score was calculated from MERS-CoV and SARS-CoV-1 binding:

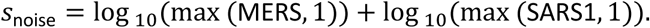

In the study population, robust true pre-existing memory against MERS-CoV or SARS-CoV-1 was considered biologically improbable. Therefore, high binding to these antigens was interpreted primarily as a marker of non-specific assay reactivity, polyreactivity, or broad low-affinity coronavirus cross-reactivity rather than antigen-specific memory.

For each endemic seasonal hCoV antigen, an ordinary least-squares regression model was fitted with the log10-transformed hCoV signal as the dependent variable and *s*_noise_as the independent variable. For example, for OC43:

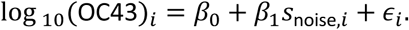

The residual from this model was extracted as the corrected hCoV titer:

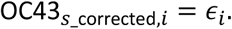

The same procedure was applied to HKU1, 229E and NL63, generating:

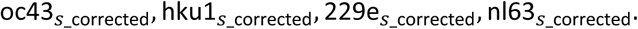

These residualized features represent the component of each endemic hCoV signal that remained after removing the portion statistically explained by broad heterologous coronavirus reactivity.

### Beta/Alpha immune-bias score

To quantify whether baseline endemic coronavirus immunity was biased toward Beta-coronaviruses or Alpha-coronaviruses, we calculated a Beta/Alpha immune-bias score. OC43 and HKU1 were treated as seasonal Beta-coronaviruses, whereas 229E and NL63 were treated as seasonal Alpha-coronaviruses. The ratio was defined as:

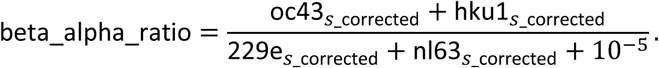

The small constant was added to avoid division by zero. This metric was designed to capture whether pre-existing coronavirus immunity was relatively enriched for OC43/HKU1-like Beta-coronavirus reactivity, which may be more structurally relevant to SARS-CoV-2 than Alpha-coronavirus reactivity.

### Endemic hCoV breadth score

A cumulative endemic hCoV breadth score was calculated to quantify the diversity of pre-existing seasonal coronavirus immunity. For each of the four endemic hCoVs, participants were assigned a binary indicator according to whether their cross-reactivity corrected residual titer exceeded the cohort median:

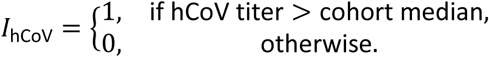

The hCoV breadth score was then calculated as the sum of the four binary indicators:

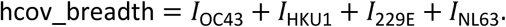

The resulting score ranged from 0 to 4, with higher values indicating broader pre-existing endemic coronavirus antibody reactivity.

### Definition of vaccine-response phenotypes

For machine-learning classification, participants were classified using an extreme-phenotype design based on the primary SARS-CoV-2 vaccine-response endpoint. Participants in the upper quartile of the response distribution were assigned to the upper-quartile response group (N=73), and participants in the lower quartile were assigned to the lower-quartile response group (N=73). Participants in the middle two quartiles were not included in the binary classification analysis. This design was used to enrich for biologically distinct response phenotypes and to test whether baseline hCoV-derived features could discriminate the extremes of vaccine responsiveness.

The primary vaccine-response endpoint used for phenotype assignment was log10 SARS-CoV-2 Spike antibody binding at v2+28. Quartile thresholds were calculated within the eligible analysis cohort.

### SIMON machine-learning analyses

Systems immunology analyses were performed using SIMON, Sequential Iterative Modeling “OverNight,” an automated machine-learning framework developed for biomedical and immunological datasets with incomplete features. SIMON has previously been described as an automated platform for comparing machine-learning approaches and identifying immune signatures in vaccine-response datasets.

Candidate predictors of SARS-CoV-2 vaccine serological response magnitude included baseline log10-transformed coronavirus serology, cross-reactivity-corrected hCoV residuals, Beta/Alpha homology-bias score, and hCoV breadth score. The primary binary outcome was Upper-quartile Responder versus Lower-quartile Responder status (N=146).

Missing predictor values were handled using nested K-nearest-neighbors imputation. To avoid information leakage, imputation parameters were estimated within the training data for each resampling split and then applied to the corresponding held-out data. Continuous predictors were standardized to zero mean and unit variance within the training data before penalized modeling.

Feature selection was performed using L1-regularized logistic regression (Lasso) embedded within a Recursive Feature Elimination with Cross-Validation (RFECV) framework. L1 regularization adds a penalty proportional to the absolute value of the model coefficients *(λ* ∑ |*βj*|) to the logistic loss function, driving the coefficients of uninformative, noisy, or redundant predictors strictly to zero. A Stratified 5-Fold cross-validation loop was utilised within RFECV to iteratively eliminate the lowest-ranked features based on area under the ROC curve, deriving the most parsimonious predictive signature.

### Permutation testing

To determine whether the observed predictive performance exceeded that expected by chance, a 1,000-iteration permutation test was performed. In each iteration, Upper-quartile Responder and Lower-quartile Responder labels were randomly permuted while the predictor matrix was held constant. The full modeling workflow was then repeated using the permuted labels. The empirical null distribution of model performance was compared with the observed model performance obtained using the true labels.

Permutation significance was calculated as:

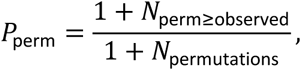

where *N*_perm≥observed_ is the number of permutations in which the model performance was equal to or greater than the observed performance. With 1,000 permutations, the minimum attainable empirical P value was 0.001.

### Longitudinal antibody-waning analysis

To assess whether baseline hCoV immune features were associated with durability of vaccine-induced antibody responses, a participant-specific antibody-decay metric was calculated using paired post-vaccination measurements. For each participant, the log10 antibody decline between approximately 28 and 180 days after the second vaccine dose was defined as:

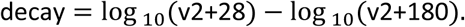

Higher values indicate greater antibody waning between the two time points. This two-timepoint contrast was used as a conservative participant-level summary measure of antibody decay, preserving one independent observation per participant and avoiding assumptions about repeated-measures covariance structure required by linear mixed-effects models.

Associations between baseline immune features and antibody decay were tested using ordinary least-squares regression. The primary models assessed the relationship between decay and hCoV-derived features, including cross-reactivity-corrected OC43 and HKU1 titers, Beta/Alpha homology-bias score, and hCoV breadth. Regression coefficients, 95% confidence intervals and two-sided P values were reported.

### Statistical analysis

Unless otherwise stated, analyses were performed on log10-transformed serological data. Ordinary least-squares regression was used for residualization of hCoV titers and for cross-sectional modeling of antibody decay. Machine-learning classification analyses were performed using nested imputation, L1-regularized feature selection, recursive feature elimination with cross-validation and permutation testing, as described above. Statistical tests were two-sided unless otherwise specified. No formal adjustment for multiple testing was applied to the machine-learning discovery analysis; instead, empirical significance was assessed using permutation testing.

## Code availability

Custom Python code used to pre-process serological data, implement stringent cross-reactivity penalization, compute Beta/Alpha homology-bias and hCoV breadth scores, perform RFECV machine-learning classification and permutation testing, and model longitudinal antibody decay is publicly available on GitHub at https://github.com/atomiclaboratory/CoV_manuscript.

## Data availability

The datasets generated and/or analysed during the current study are available on Zenodo.

## Acknowledgements

This research was supported by a Coalition for Epidemic Preparedness Innovations (CEPI) grant aiming to determine SARS-CoV-2 correlates of protection.

## Author information

Authors and affiliation

**Centre for Human Genetics and Pandemic Sciences Institute, Nuffield Department of Medicine, University of Oxford, Oxford, United Kingdom**

Stephanie Longet, Caolann Brady, Tom Tipton & Miles Carroll

**Pandemic Sciences Institute, University of Oxford, Oxford, United Kingdom**

Stephanie Longet, Caolann Brady, Tom Tipton & Miles Carroll

**International Center for Infectiology Research (CIRI), Team GIMAP, Claude Bernard Lyon 1 University, Inserm, U1111, CNRS, UMR530, Saint-Etienne, France**

Stephanie Longet & Salma Bessalah

**Nuffield Department of Clinical Medicine, Peter Medawar Building for Pathogen Research, University of Oxford, Oxford, United Kingdom**

Eleanor Barnes, Susanna Dunachie & Paul Klenerman

**Oxford University Hospitals NHS Foundation Trust, Oxford, United Kingdom.**

Eleanor Barnes, Susanna Dunachie & Paul Klenerman

**Nuffield Department of Clinical Medicine, Oxford Centre For Global Health Research, University of Oxford, Oxford, United Kingdom**

Susanna Dunachie

**Mahidol-Oxford Tropical Medicine Research Unit, Mahidol University, Bangkok, Thailand**

Susanna Dunachie

**Translational and Clinical Research Institute Immunity and Inflammation Theme, Newcastle University, Newcastle, United Kingdom**

Christopher JA Duncan

**Department of Infection and Tropical Medicine, Newcastle upon Tyne Hospitals NHS Foundation Trust, Newcastle, United Kingdom**

Christopher JA Duncan

**Translational Gastroenterology Unit, University of Oxford, Oxford, United Kingdom**

Paul Klenerman

**NIHR Oxford Biomedical Research Centre, University of Oxford, Oxford, United Kingdom**

Paul Klenerman

**Institute of Cancer and Genomic Science, College of Medical and Dental Science, University of Birmingham, Birmingham, United Kingdom**

Alexa Richter

**University Hospitals Birmingham NHS Foundation Trust, Birmingham, United Kingdom**

Alexa Richter

**MRC Toxicology Unit, University of Cambridge, Cambridge, UK**

James ED Thaventhiran

**NIHR Health Protection Research Unit in Emerging and Zoonotic Infections, Institute of Infection, Veterinary and Ecological Sciences, University of Liverpool, Liverpool, United Kingdom.**

Lance Turtle

**Liverpool University Hospitals NHS Foundation Trust, Liverpool, United Kingdom.**

Lance Turtle

**Department of Infection, Immunity and Cardiovascular Disease, University of Sheffield, Sheffield, United Kingdom.**

Thushan I de Silva

**Sheffield Teaching Hospitals NHS Foundation Trust, Sheffield, United Kingdom.**

Thushan I de Silva

**National Emerging Infectious Diseases Laboratories, Boston University, National Emerging Infectious Diseases Laboratories, 620 Albany St, Boston, MA 02703, United States**

Ivan Tomic & Adriana Tomic

**Department of Virology, Immunology & Microbiology, Boston University Medical School; Boston, MA, US**

Ivan Tomic & Adriana Tomic

**Biomedical Engineering, Boston University, College of Engineering; Boston, MA, US**

Ivan Tomic & Adriana Tomic

## Contribution

S.L., C.B.: MSD assays in PITCH studies, pre-processing of data and writing original draft. I.T.: conceptualization and development of the systems immunology methodology, formal analysis, software/implementation, data curation for analytical feature engineering, development of the SIMON analytical strategy, running SIMON analyses, machine-learning and longitudinal modeling, visualization/interpretation of results, and writing original draft sections. T.T.: MSD assays in PITCH studies. S.B.: pre-processing of data. E.B.: review and editing. PI of Oxford PITCH cohort; S.D.: review and editing. PI of Oxford PITCH cohort; C.D.: review and editing. PI of Newcastle PITCH cohort; P.K.: review and editing. PI of Oxford PITCH cohort; A.R.: review and editing. PI of Birmingham PITCH cohort; J.T.: review and editing. PI of Cambridge PITCH cohort; L.T.: review and editing. PI of Liverpool PITCH cohort; TdS.: review and editing. PI of Sheffield PITCH cohort; A.T.: conceptualization of systems immunology methodology, oversight of SIMON analyses, review and editing. M.C: Conceptualization, funding acquisition, methodology, project administration, supervision, review, and editing.

## Ethics declarations

Author S.L. serves as an Associate Editor of this journal and had no role in the peer-review or decision to publish this manuscript.

Author M.C. has previously received free MSD reagents worth approximately £20k. However, these reagents were not used for this study and MSD played no role in this project.

All other authors declare no financial or non-financial competing interests.

